# Influence of different anionic charges in lead compoundsin the remediation of lead by*Eleusine indica* (Gaertn)

**DOI:** 10.1101/2021.11.22.469534

**Authors:** B. Ikhajiagbe, R. I. Ehiarinmwian, G.O. Omoregie

## Abstract

The study was carried out to investigate the remediative capacity of *Eleusine indica* in lead-polluted soil. Soil samples were collected near student hostel (hall 5) in the University of Benin. The soil samples were sun dried to constant weight and was pulverized with wooden roller and sieved with a hand sieve of 2 mm mesh size. The sieved soil was spiked with 0.625 g lead nitrate (PbNO_3_), lead sulphate (PbSO_4_), lead carbonate (PbCO_3_), lead acetetrahydrate (PbC_2_H_6_) and lead chloride (PbCl_2_) separately in three replicate using aqueous standard solutions. Tillers of Eleusine Indica were placed in the metal – polluted soil immediately and the experiment was allowed to stay for 15 weeks. The result showed that the uptake efficiency for *Eleusineindica* in both shoots and roots for lead nitrate was 0.016% and 0.8%, lead sulphate 0.016 % and 0.352 %, lead carbonate 0.064% and 0.496 %, lead acetetrahydrate 0.032 % and 0.688 %, and lead chloride 0.08 % and 0.72 % respectively, indicating that the plant might have sequestered the metal in the soil rather than accumulating it in the leaves. This was evident in the presentation of the metal sequestration factor of over 70 % irrespective of the nature of the metal. Microbial count of soil before and after contamination with lead nitrate was 1.9×104 and 0.4×104 cfu/g indicating a reduction. The study therefore revealed that *Eleusine indica* is a high efficient plant in sequestering lead in polluted soil.

## INTRODUCTION

The environment has always been under natural stress but its degradation was not as severe as it is today. The pollution of the environment by heavy metals is now a world issue requiring considerable attention. Soils polluted with heavy metals often lack established vegetation cover due to the harmful effects of the heavy metal or incessant physical disturbances such as erosion (Alfaraas*et al*., 2014). Most heavy metal emission results from anthropogenic sources such as transportation and industries. Herbicides, manures and sewage silt used in agriculture are also sources of heavy metals in the environment (Alkorta*et al*., 2004). The persistence of these heavy metals in soils as well as continuous exposure to them can indirectly or directly lead to their buildup in plants, animals and eventually humans. The present study emphasizes the effects of heavy metals.

Heavy metals (HM) can be defined as non-biodegradable persistent inorganic chemical components with the atomic mass over 20 amu and density greater than 5g/cm^−3^. They have cytotoxic and mutagenic effect on humans, plants and animals (Anoliefo*et al*., 2006; Ikhajiagbe and Anoliefo, 2011). Coincidentally there is no clear suggestion of what a heavy metal is, density is used at times to be the defining factor (Hossain *et al*., 2012).

Heavy metals have become a worldwide problem in soil pollution bringing about losses in agricultural yield as well as hazardous health effects as they enter the food chain (Aiyesanmi*et al*., 2012). As a consequence of modern land-use techniques, or activities that mobilize heavy metals, many soils are polluted by different agents such as heavy metals, nutrients and pesticides. Rapid urbanization, use of pesticide and mineral fertilizer have led to toxic metal pollution of land, plant, as well as water resources (Flora and Pachauri, 2010). This could be credited to the use of polluted irrigation water, the addition of some herbicides, and also pollution from traffic. Sewage sludge, agricultural manures, and sea bird manure in natural ecosystems, industrial activities, fuel, and automobile tyres can also be significant metal sources (Ikhajiagbe*et al*., 2012). Of specific note, mining operations and coal burning usually increases the levels of heavy metal as studied by Emoghene and Futughe (2016). Heavy metals can amass in plants via both root and foliage systems. Many research have been conducted to ascertain the harmful levels of heavy metals for certain plants, particularly those metals regarded as public health threats. Heavy metal toxicity to plant metabolism has received broad research interest for some decades (Ezekiel, 2015). This is dire in soils where agricultural production thrives, because contaminants can be amassed in crops. This can lead to human health problems.

The existence of heavy metals in the soil arises from weathering of the parent constituents at intensities that are detected as trace (<1000mgkg^−1^) and seldom toxic (Green, 2011). Due to human interference with nature’s natural resources most soils might accumulate heavy metals beyond tolerable values, this can affect human wellbeing, plants, animals, ecosystems, or further media.

Lead (Pb) which is well known has been in existence since ancient times. It is naturally available in the earth’s crust in small amounts, but for centuries it has been mined and distributed all over the environments from where it has slowly become integrated into the structural tissue of plants, animals and humans (Jiang and Liu, 2010). Lead exists as Pb^+2^ion during the chemical reaction. Lead is known as a highly toxic metal as well as a cumulative poison. The industries often dealing with lead are the battery manufacturing, cable making, motor vehicle repair and metal grinding industries. Lead is also used in piping, accumulators, conducting materials, lead chambers, soldering, anti-knock substances, printing characters, colored pigments, wrappings for food, tobacco, radiation shielding and as an additive in gasoline (Kavita*et al*., 2014). Therefore this study was carried out to evaluate the to investigate the remediative capacity of *Eleusine indica* in a lead-polluted soil.

## MATERIALS AND METHODS

### Study Area

The site chosen for the research was a space in the general laboratory close to the Botanic Garden, Ugbowo campus of University of Benin, Benin City, Nigeria, which lies within the rainforest ecological zone of Midwestern Nigeria.

### Collection and Preparation of Materials for the Experiment

Soil used in the present study was collected from an area measuring 50 m x 50 m marked near hall 5 in university of Benin. Top soil (0-10 cm) was collected randomly from the marked plot. Thereafter, 10 kg sun-dried soil each was placed into non perforated bowls measuring (14.6 cm height and diameter of 22.4 cm).

Measured quantity of soil placed in each bowl occupied a dimension of 10.4 cm in depth and a radius of 13.2 cm. surface area of top soil thus measured was 891.6 cm^2^.

### Procurement and Subsequent Application of Lead Metal

Prior to conduction of field experiment, the researcher went out on a survey to determine the choice of metal to be used for the research. Lead metal was later concluded to be used for the experiment due its availability and negative impact on soil, plant and the environment at large.

Soils in each bowl were wetted with the test lead metal. 0.625 g of lead metal was dissolved in 250 ml of water that was subsequently placed in the non-perforated bows.

There were 6 different levels of pollution: lead carbonate, lead chloride, lead nitrate, lead sulphate, lead acetetrahydrate and the control. The lead metal was dissolved in 250ml of water. The control was wetted with ordinary water. The concentrations were obtained as follows:

A. 0.625g lead nitrate + 250 ml water used to wet 10 kg of soil
B. 0.625g lead sulphate+ 250 ml water used to wet 10 kg of soil
C. 0.625g lead carbonate+ 250 ml water used to wet 10 kg of soil
D. 0.625g Lead acetetrahydrate+ 250 ml water used to wet 10 kg of soil
E. 0.625g lead chloride + 250 ml water used to wet 10 kg of soil
F. 250 ml water used to wet 10 kg of soil. No metal added.

These treatments were replicated 3 times in completely randomized block design. The entire set up was left to naturally attenuate for some hours before tillers of *Eleusine indica*, the test plant, were transplanted into it.

### Nursery for *Eleusine indica*

Prior to the commencement of the study, fresh plants, showing no evidence of discoloration or morphological distortion, were selected from an open field in the Faculty of Agriculture, University of Benin, Benin City. The plant was immediately divided into equal tillers. These were sown in already prepared nursery, which consisted of different bowls as used in the study in other to accommodate more Eleusine. After a month, freshly emerged tillers of similar size and height were harvested and used as transplants in the study, as reported earlier.

### Parameters Considered

The experimental set up, consisting of lead heavy metal-polluted soils and the transplanted test plants, were left close to the windows of the general laboratory to avoid rain penetration as the metal was to be total conserved without run off. For additional 3 months. (March through June, 2018).Paramters determined during the course of the study have been indicated (Table 1).

**Table 1:** Morphological Parameters Accessed

| S/N | Parameter | Method of determination |
| --- | --- | --- |
| 1 | Plant Height (cm): | By tape rule for measurement |
| 2 | Stem Width (mm): | By Vernier calliper |
| 3 | Flag leave blade length (cm): | By tape rule |
| 4 | Flag leave blade width (mm): | Vernier calliper |
| 5 | No of additional spike: | By physical counting |
| 6 | Peduncle length: | tape rule for measurement |
| 7 | Length of longest spike (cm): | tape rule for measurement |
| 8 | Length of spikelet (mm): | tape rule for measurement |
| 9 | Number of spikelet's per plant: | physical counting |
| 10 | Depth of longest root: | tape rule for measurement after plant had been uprooted, washed and dried |
| 11 | Internode length (cm): | tape rule for measurement |
| 12 | No. of Culm branching per plant: | physical counting |
| 13 | No. of tillers per plant: | physical counting |
| 14 | Plant dry wt. (g): | Determined by first uprooting the plant, and then carefully washing the sand particles from the roots. The entire plant (shoot and roots altogether) was dried in the oven at 35°C for 30 minutes. The dried plant was weighed in an electric balance. |
| 15 | Plant moisture content (g): | The difference between the plant fresh weight and plant dry weight was given as the moisture content in grammes/plant |
| 16 | No of main root: | Determined by physical counting after plant had been uprooted and the roots carefully washed and dried. |
| 17 | No of Branch root: | physical counting after plant had been uprooted and the roots carefully washed and dried. |
| 18 | First day to flowering (DAT): | physical counting and observation, taking the dates into note |
| 19 | No of leaves: | physical counting |
| 20 | Number of spike: | physical counting |
| 21 | Total No of primary root branches: | physical counting after plant had been uprooted and the roots carefully washed and dried. |
| 22 | Percentage chlorotic plant population: | This was determined as the percentage of the total plants per bowl that showed significant signs of leaf chlorosis. For the purpose of this study, plants that had up to >10 % of its entire foliage being chlorotic were defined as chlorotic plants. Therefore, the percentage of these chlorotic plants over the entire plant population per treatment was defined in the study as Percentage chlorotic plant population |

### Soil Microbial Analysis

#### Collection of Bulk and Rhizospheric Soil

The plant was carefully uprooted and shaken carefully for some minutes to get rid of the large soil particles. The rhizospheric soil were the ones directly attached to the roots of the plant (< 3 mm thick). The soils were collected by using a brush to carefully remove the soils so that the morphological presentations of the roots were not compromised. The soils collected were pooled for each treatment, wrapped in aluminum foil paper and taken to the laboratory for analysis.

The bulk soil collected consisted of those soils collected that had no direct association with the root of the plant and this was done to the control bowls alone. These were collected within a radius of > 15 cm from the base of each plant and at a 10 cm depth. Soils collected were also pooled for each treatment, wrapped in aluminum foil paper and taken to the laboratory for analysis.

#### Preparation of Soil Sample Dilution Series

The soil sample were air-dried for 72 hrs and was sieved to remove undesirable material. The dilution series for soil sample was done by transferring 1g of soil sample to 9ml of normal saline in sterile test tube as blank. The test tubes were mechanically shaken for about 30 seconds to get homogenous mixture and were taken as 10^−1^ dilution factor, 10ml were transferred from the 10^−1^dilution into another 9ml blank to obtain a 10^−2^ dilution and same process was repeated five times to obtain a dilution factor of 10^−7^.

A tablet of Ketacon (200 mg) was grinded and 2 capsules of chloramphenicol (250 mg) were dissolved in the nutrient agar and potato dextrose agar to support the bacterial and fungal growth.

#### Heterotrophic Bacterial and Fungal Counts

The pour plate method was employed in taking the bacterial counts. Serially diluted portion of the 10^−7^ of each soil sample for both bulk and rhizopheric was inoculated into the nutrient agar plates for bacterial and potato dextrose agar plates for fungal counts. The plates were inoculated at room temperature for 72 hrs respectively, for bacterial and fungal growth. After incubation, colonies were then counted and colony forming unit (cfu/g) of the soil samples was determined.

#### Determination of Lead (Pb) From Shoot, Root, and Soil Samples

Firstly the samples where oven dried at 100 degree Celsius and was crushed into a smaller particle size and1g was weighed into kjedahl flask and 10ml of a mixed acid (HCLO_4_ and HCL 1:1) was introduce and then heated for 45 minutes in the fumes chamber until the brown fume gradually disappear and it became clear. The digest was filtered using whatman Number 1 filter paper, and the filtrate was made to mark. Bulk scientific, VGP210, Atomic absorption spectrophotometer was used to analyze for Pb. Hollow cathode lamp used for absorbance measurements. Wavelength 283.5, standards1, 2, 3, 5, 10 ppm, slit 7 where used respectively.

## RESULTS

Prior to sowing of tillers of *E. indica*, morphological parameters of the plant were recorded and presented on Table 4.1. Average plant height of the tillers used was 16.3 cm with a flag leaf blade length of 14.2 cm. There were at least 4 leaves per culm and there were no evidences of chlorosis and necrosis.

Plant height of *E. indica* presented within 15 weeks following exposure to test metal showed progressive increases with time (Fig. 1). No statistical differences in the heights of test plant exposed to lead nitrate, chloride, carbonate, acetetrahydrate as well as sulphate, was recorded at each week. Plant height ranged from 21.2 – 25.5 cm in the metal-exposed plants compared to 20.5 cm in the control at the 3^rd^ week, whereas plant height at the 15^th^ week ranged from 28.4 – 32.4 cm irrespective of treatment applied or control.

**Figure 1:**
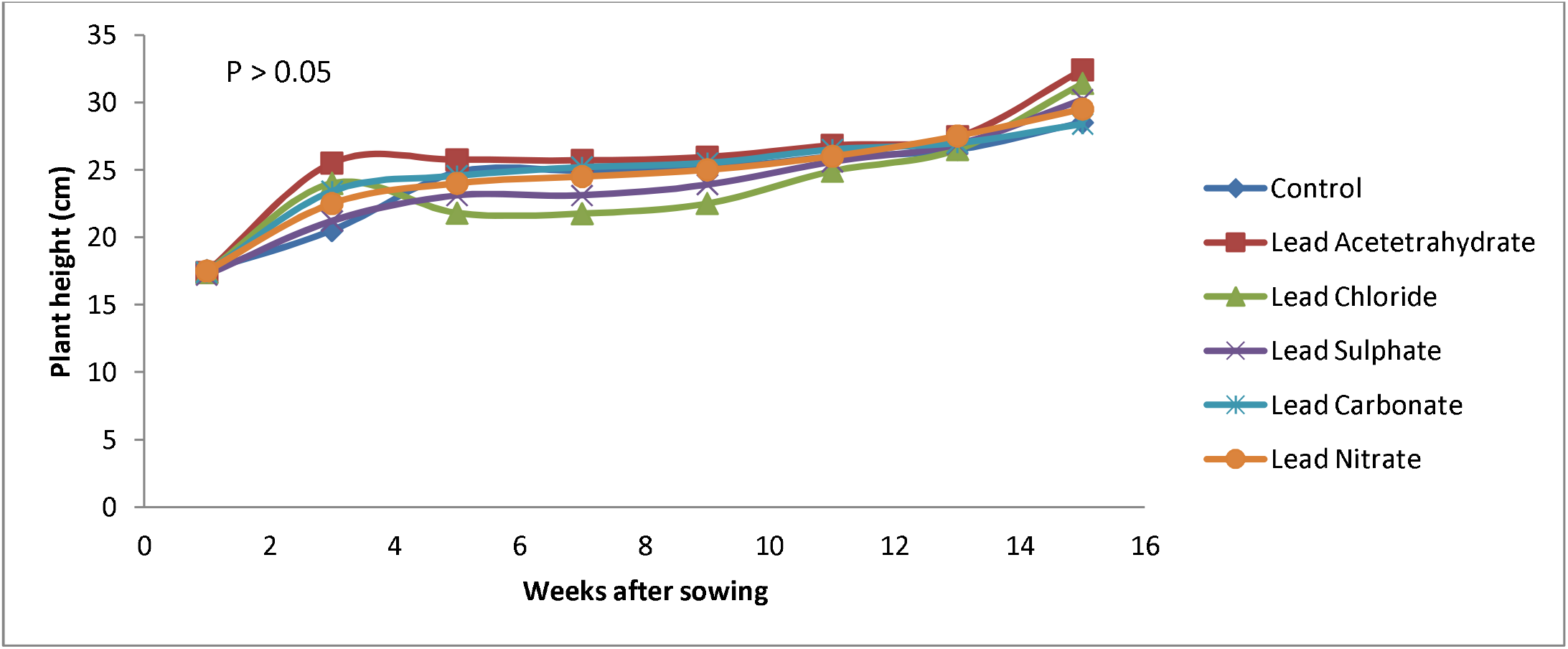
Plant height of *E. indica* weeks after sowing.

Similarly, stem width progressively increased with time, but show minimal differences among the various treatments, with values ranging from 2.96 – 3.55 mm at the 3^rd^ week after transplanting the tillers (Fig. 2). At the 15^th^ week, stem width generally fell within the range of 4.80 mm – 4.96 mm.

**Figure 2:**
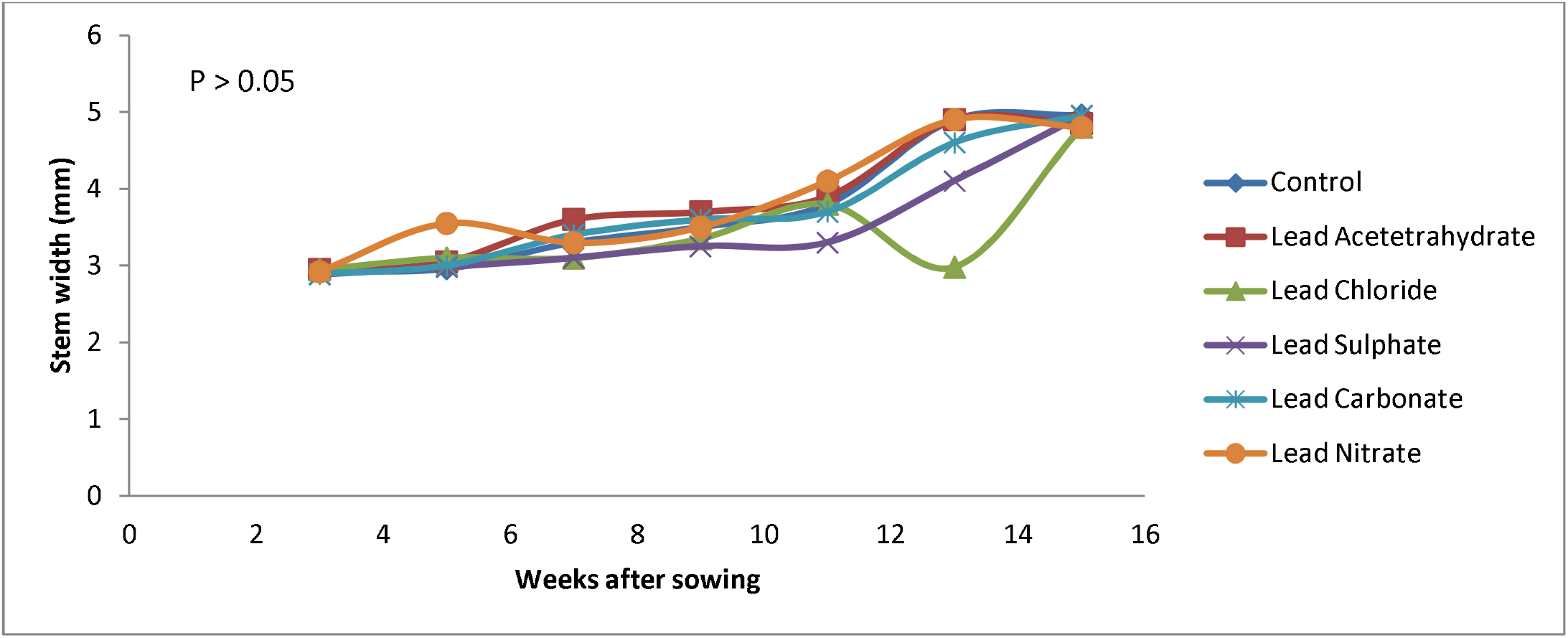
Stem width of *E. indica* weeks after sowing

Progression in plant leaf blade length between the 1^st^ and 15^th^ week after sowing was monitored (Figure 3). At 3^rd^ week after sowing, leaf blade length ranged from 18.58 cm in lead carbonate-impacted soil, compared to 25 cm in lead acetetrahydrate. Expectedly comparable observation was noted for leaf blade breath of lead chloride-exposed plants at the 7^th^ week (19.54 cm – 24.1) and 28.0 – 31.54 cm) at the 15^th^ week. Generally, throughout this period, there was no significant difference in the leaf blade length of plant under the present experimental condition.

**Figure 3:**
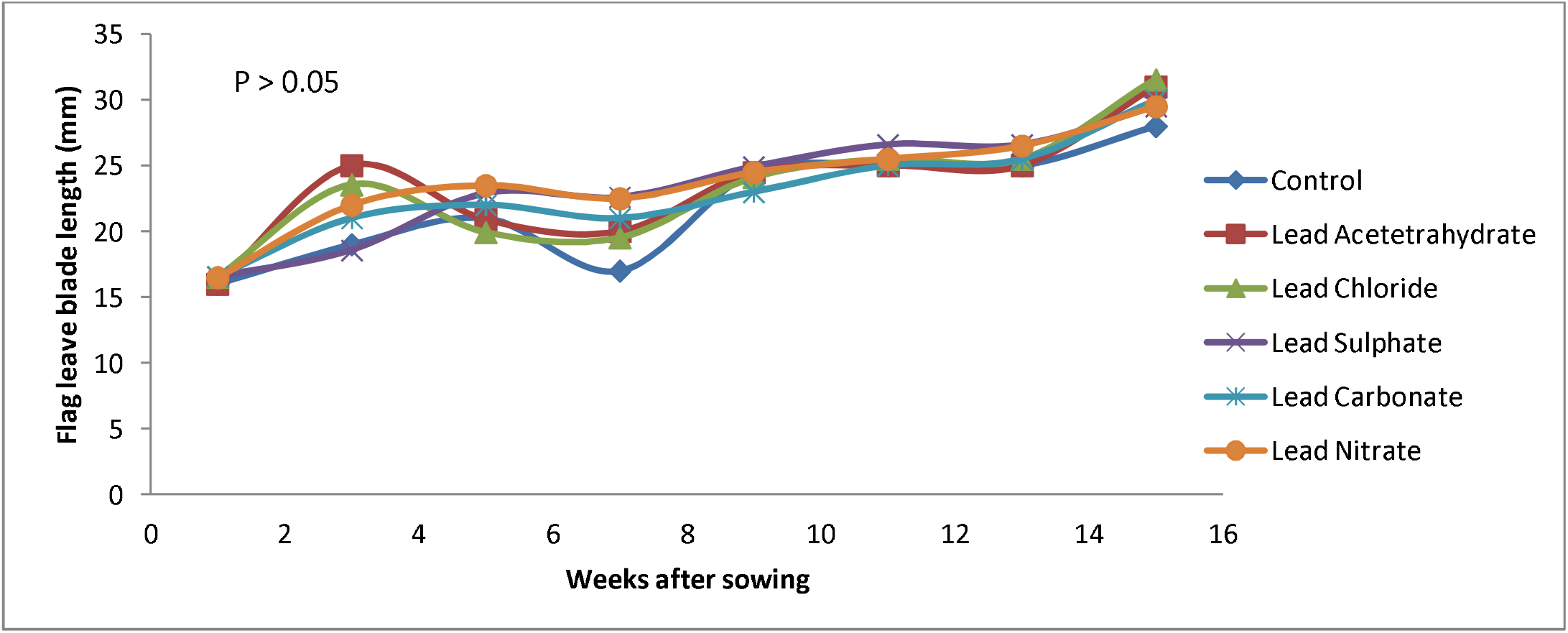
Flag leave blade length of *E. indica* weeks after sowing

The development in plant flag leave blade width under experimental influence (figure 4). At week 3internode ranged from 8 cm in lead nitrate-exposed plants to 9 cm in lead chloride-treated plants, and 8cm in the control. This generally indicated that there was no significant difference in flag leave blade width among the plants under experimental condition from week one of the experiments to week 15. Also no sharp difference in the internodes of all the lead metal was recorded.

**Figure 4:**
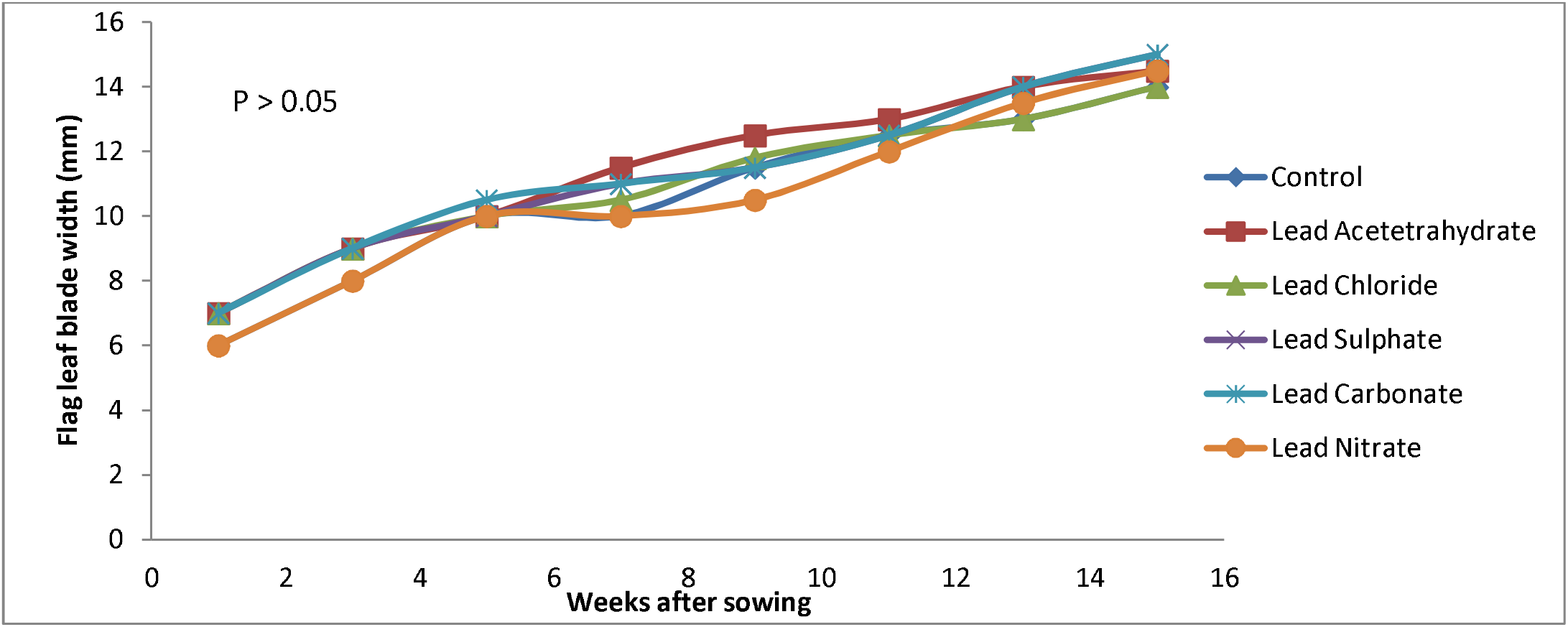
Flag leave blade width of *E. indica* weeks after sowing

Compared to the control (28.5 cm), plant height in the metal-exposed soils did not significantly differ from the former (28.4 – 32.4 cm) at the 15^th^ week following transplanting of test plant (p>0.05) (Table 2). Similarly, no differences were reported for stem width (4.80 – 4.96 mm) between control and metal-impacted plants. There were generally no significant changes in plant morphology when exposed to the various presentations of lead metal, and compared to the control. However, there were more newly emerging species of plant species within the surrounding of the control plant than around the metal-exposed plant.

**Table 2:**
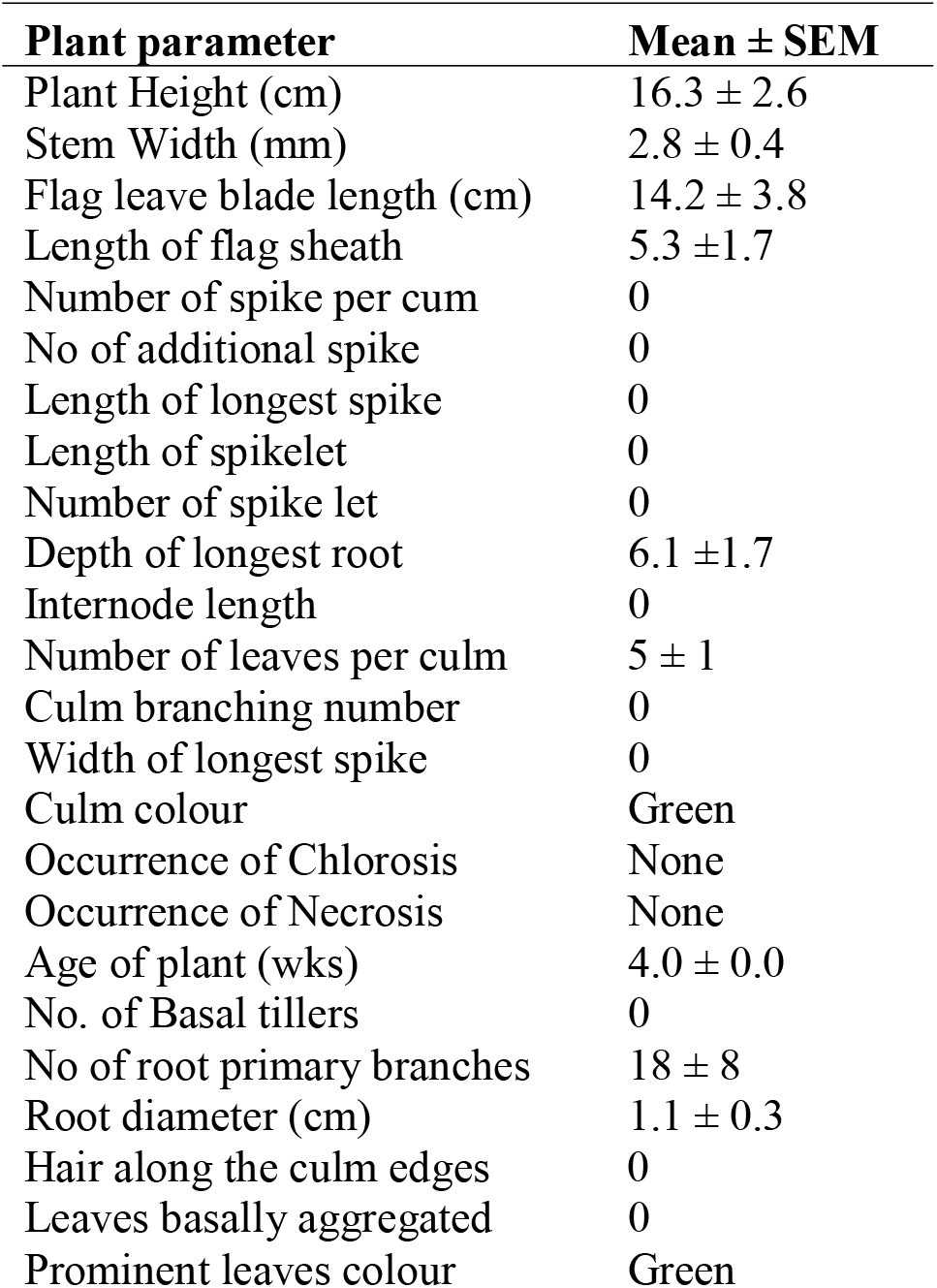
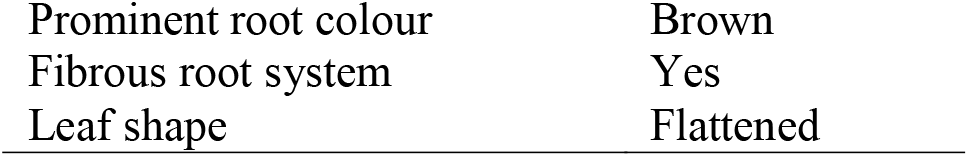
Plant parameters before transplanting

**Table 2:**
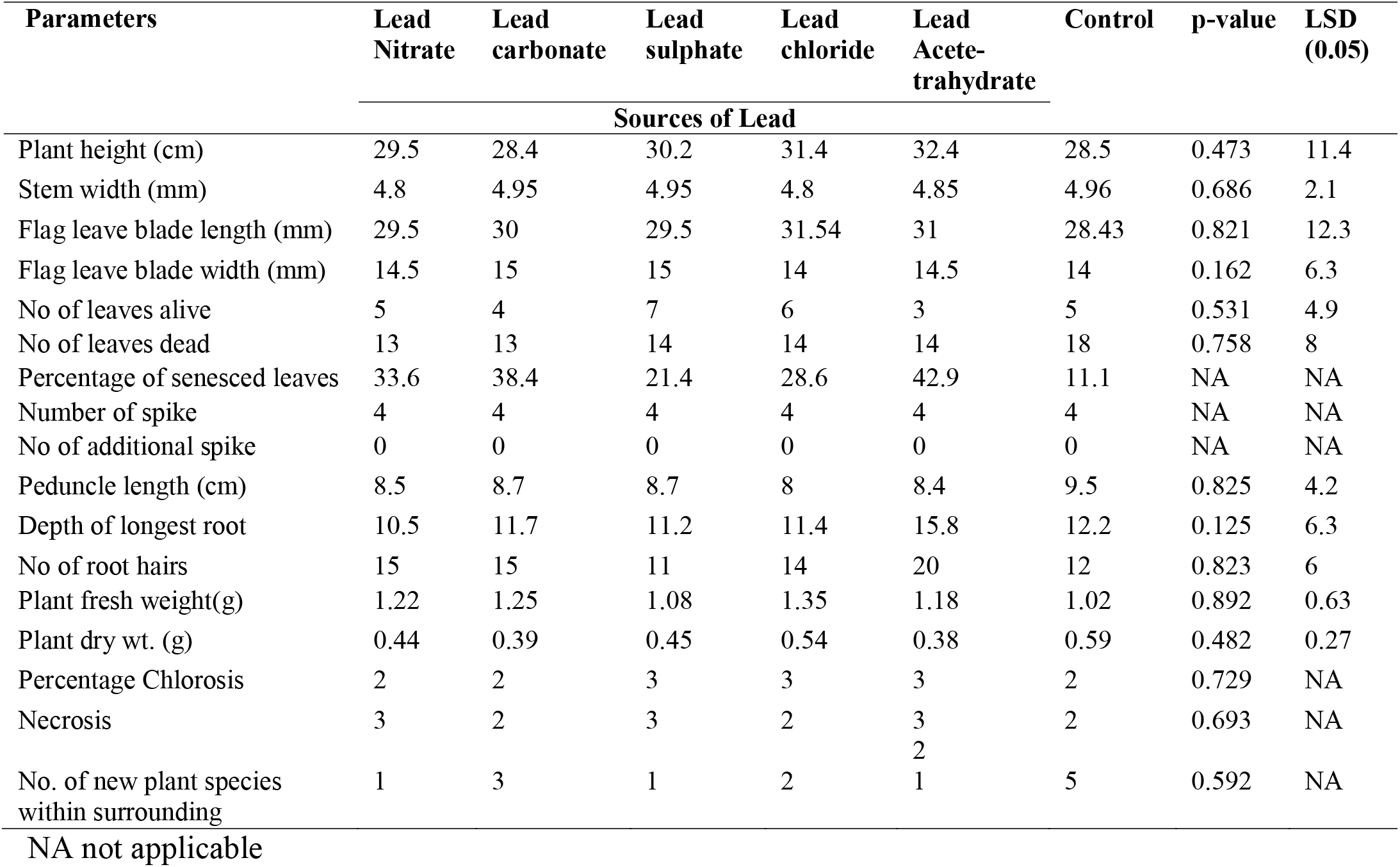
Morphological parameters at 15 weeks after sowing and exposure to Pb.

The accumulation capacity of the test plants is shown in Table 3. For metal accumulation in shoot, there were no statistical differences in outcome (0.01 – 0.05 mg/kg). Similarly, accumulation of metal in the roots followed similar pattern (0.22 – 0.50 mg/kg). Remediation efficiency within shoot and root were generally below 0.80%.

**Table 3:** Accumulated Pb in *E. indica* plant

|  | Accumulated values (mg/kg) |  | Uptake efficiency (%) |  |
| --- | --- | --- | --- | --- |
|  | Shoot | Root | Shoot | Root |
| Lead Nitrate | 0.01 | 0.5 | 0.016 | 0.8 |
| Lead carbonate | 0.01 | 0.22 | 0.016 | 0.352 |
| Lead sulphate | 0.04 | 0.31 | 0.064 | 0.496 |
| Lead chloride | 0.02 | 0.43 | 0.032 | 0.688 |
| Lead Acetate tetrahydrate<br>(Start = 62.5 mg/kg) | 0.05 | 0.45 | 0.08 | 0.72 |
| p-value | 0.517 | 0.361 | NA | NA |

NA not applicable

Residual heavy metal concentration in soil sown with test plant has been presented on Table 4. As presented, initial soil concentration was 62.5 mg/kg, but residual concentration after plant exposure for 15 weeks was 19.55 mg/kg which represented a remediation efficiency of 68.39%. The most reduced metal concentration in soil was with lead sulphate (4.05 mg/kg), an efficiency of 93.07%. Similarly, total sequestered Pb was highest when lead type in soil was sulphate (58.1 mg/kg) than when other lead anions were present (Table 4). This implied that the contaminant sequestration factor was 92.96%. Generally, sequestration factor ranged from 67.9 – 92.96%. This means that the plant was most likely to sequester the metal than bioaccumulate it in the above ground plant parts.

**Table 4:** Residual heavy metal concentration in soil sown with test plant

|  | Residual Pb |  | Sequestered Pb |  |
| --- | --- | --- | --- | --- |
|  | Residual Soil conc. (mg/kg) | Remediation efficiency | Total Sequestered Pb (mg/kg) | Sequestration factor (%) |
| Lead Nitrate | 19.55 | 68.39 | 42.44 | 67.9 |
| Lead carbonate | 16 | 74.04 | 46.27 | 74.03 |
| Lead sulphate | 4.05 | 93.07 | 58.1 | 92.96 |
| Lead chloride | 6.65 | 88.93 | 55.4 | 88.64 |
| Lead Acetate tetrahydrate<br>(Start = 62.5 mg/kg) | 11 | 82.01 | 51 | 81.6 |
| p-value | 0.012 | NA | 0.363 | NA |

NA not applicable

Quantitative bacteria composition of the soil before and after plant exposure has been presented (Table 5). There was a 78.94% decrease in rhizospheric bacterial population in soils polluted with lead nitrate, compared to that polluted with lead carbonate (6.81% reduction). However, growth enhancing effect of metal on rhizospheric bacterial population was reported in soil impacted with the sulphate, chloride and acetetrahydrate of the metal respectively. Bacteria population increased by as high as 200% of initial 0.4× 10^4^cfu/g concentration 15 weeks earlier when soil was polluted with the acetetrahydrate of the metal.

**Table 5:** Quantitative composition of culturable rhizospheric bacteria during and after plant exposure.

| | Before contamination<br>($\times 10^4$cfu/g) | 15 weeks after contamination<br>($\times 10^4$cfu/g) | p-value | Percentage change (%) / remark |
| --- | --- | --- | --- | --- |
| Lead Nitrate | 1.9 | 0.4 | 0.031 | 78.94/ reduction |
| Lead carbonate | 0.88 | 0.82 | 0.836 | 6.81 / reduction |
| Lead sulphate | 0.2 | 0.3 | 0.637 | 50.00/ increase |
| Lead chloride | 0.5 | 0.6 | 0.701 | 20.00/ increase |
| Lead Acetate tetrahydrate | 0.4 | 1.2 | 0.021 | 200.00/ increase |
| Control | 0.5 | 0.4 | 0.832 | 20.00/ reduction |
| (Rhizospheric soil) |  |  |  |  |
| Control | 0.1 | 1.3 | 0.041 | 1200.00/ increase |
| (Bulk soil) |  |  |  |  |
| p-value | 0.031 | 0.042 | NA | NA |

## DISCUSSION

### Morphological Development of the Plant

In this study, it was recorded that at 15 weeks after transplanting the stem width of *Eleusine indica* increased from 2.8 mm to 4.96 mm, a range of 4.8 to 4.9 mm in all test exposed to Pb treatment. Significant increase in plant height was recorded in all test plants. Previous studies support the idea that plant physiology is controlled by cellular activities (Krämer, 2010). The test plant in the control experiment showed similar increase in stem width and plant height and this buttress the fact that plants exposed to Pb^2+^ had comparable morphology with unexposed plants. The root system of *Eleusine indica* not only provides an enormous surface area that absorbs and accumulates the water and nutrients that are essential for growth, but also absorbs other non-essential contaminants such as Pb and other heavy metals. However, based on the observed features of test plants in the present study, it could be that Pb was sequestered by the test plant (Lamhamdi*et al*., 2011).

Metals were found at high levels in the root than the shoot with no sign of toxicity and this would have effect on the plant height and also on the shoot of the test plant. The grass *E. indica* in this study expressed high level of Pb in its roots. One of the mechanisms by which uptake of metal occurs in the roots may include binding of the positively charged toxic metal ions to negative charges in the cell wall (Murali*et al*., 2014) and the low transport of heavy metal to shoots may be due to saturation of root metal uptake, when internal metal concentrations are high. And also this would have considerable effect on the test plant.

Number of survived leaf ranged from (5 - 7) in the exposed test plants as compared to 5 in the control. Studies have shown that Pb on plants could either be positive or negative depending on the concentrations. The increase number of leaves might be as a result of the cellular metabolic activities occurring within the cell of the test plant. Specific composition and complex interplay of a multitude of regulatory pathways of the cell of the plant must have had influence on the test plant. Such that the contaminated plant adjust to tolerate the stress, hence increase the number of leaves and shed off older leaves so as to reduce the amount of heavy metal in the plant system. This supports the findings of Nazarian*et al*. (2016) who also suggested that contamination and perturbations in the environment can have significant effect on leave number.

In terms of high root number, liberal growth was observed in exposed plant. This was confirmed by Omoregbee*et al*. (2015) who reported an increase in the number of root in *Eleusine* plants, and stated that it was as a result of the high root biomass which translate soil microorganism and their biodegradative activity through increase in diffusion, mass flow and concentration of nutrients, thus facilitating the degradation of oil hydrocarbon and other heavy metals such as lead. This is supported by reports of Anoliefo and Ikhajiagbe (2011). The increase in the morphological parameters has been directly interrelated with the increase in photosynthetic activity of the plant.

### Progression of Stress Responses in test Plant

Browning spot at leaf blade particularly of young leaves of plant. Though conclusive, this may be a natural phenomenon or of other environmental conditions not covered by the scope of this research. However, Opeolu*et al*. (2010) reported dark brown spots to be caused by manganese oxide precipitate as a result of sensitive genotype accumulating manganese in localized areas in the leaf. Amorphous spots on the leaves of *Ocimumbasilicum* L. due to the presence of cadmium has also been reported by Orisakwe*et al*. (2012). The plant may have had partial closure of stomata, which in turn result in less water movement from roots up the plant. This created an internal water deficit followed by reduction of CO_2_ uptake and photosynthesis causing stress and nutrient uptake deficiencies.

Plant wilting was generally noticed in the plants exposed to Pb metal. Pourrut*et al*. (2011) reported that wilting can actually be caused by chemical injury such as pesticide and other heavy metal. In the present study, all the bowls with Pb Impacted Soil (PIS) experienced wilting as started in Table 2 and this started with folding of plant leaves. Plants in the control showed no evidence of wilting.

Although the soils on which plants were sown showed evidence of adequate water, insufficient available soil moisture causes stresses that can lead to wilting and premature defoliation under extreme conditions. Under less extreme conditions, the stomata close to decrease the rate of transpiration. When this occurs, carbon dioxide can no longer enter into the leaves through the stomata and photosynthesis decreases. These can lead to death of the plant as stated in Table 2. Among environmental factors, water availability is probably the most limiting for crop quality and productivity, comprising economic output and human food supply (Patel*et al*., 2010). Water deficit is a multidimensional stress affecting plants at various levels of their organization (Alfaraas*et al*., 2014). Thus, the effects of stress are often manifested at morpho-physiological, biochemical and molecular level, such as inhibition of growth (Alkorta*et al*., 2004) as have been shown in Table 2, as well as changes in protein contents, among others. Some of these responses are directly triggered by the changing water status of the tissues (Hossain*et al*.,2012).

Patterns of chemical injury on individual plants differ depending primarily on whether the chemical caused damage directly on contact or is absorbed and moves throughout the plant. Wilting, as reported in the study, may have resulted from root damage. Aiyesanmi*et al*. (2012) reported that some toxic contact chemicals in the root zone, result in poor root development. Symptoms from root-contact chemicals are localized where the chemical contacts the root, but produce general symptoms in the shoot. The shoots may show water and nutrient stress symptoms, i.e. reduced growth and wilting as reported in the study. Roots are injured and root tips may be killed. This will result in a general stunted growth in the plant. Flora and Pachauri (2010) also recorded that in severe cases, wilting occurs even though the soil is wet.

Lead polluted soil exposed to the test plants showed significant increase in the number of chlorotic as well as necrotic leaves when compared to the control. Obviously, chlorosis and necrosis are indications of lead contamination whose incidences increase with metal concentration in the soil. Emoghene and Futughe (2016) also reported leaf chlorosis and necrosis in Paddy plant due to the application of lead while Ezekiel (2015) reported induced deficiency of some ions with the same electrical charge or ion radius such as magnesium and iron to be responsible for leaf chlorosis. Regions of necrotic lesions have also been reported in this study. In the control, necrosis appeared late and progressed inwards the leaf until the entire leaf tissue died out with regions of necrosis.

### Plant Productivity and Growth Indices

Chlorophyll content of *Eleusine indica* which was diligently followed up for the 11wks of the experiment was observed using morphological colour codes that ranges from (1 - 8) and this differed significantly from (green yellow - dark green) all throught the experimental test bowls. The control had between (7 green – 8 darkgreen) and this might be as a result of the fact that plants in control experiment thrived under normal growth conditions devoid of abiotic stress. Although there was a strong recovery of test plant.

Plants root traits (root length and root dry matter) are presented in table two at fifteenth week. *Eleusine indica L* showed higher root length (15 – 20 cm). Similar to the root length measurement, the root dry matter of the test plant was 0.38 – 0.54. The known hyper accumulator plants tested in this experiment showed a different root length measurement at lead acetetetrahydrate and a lower root length compare to *lead chloride*.(Table 3).Hyperaccumulator plants are characterised by: (i) tolerant to high concentration of metal in soil, (ii) accumulate/absorb metal from the soil, (iii) rapid growth rate, (iv) producing high biomass, and (v) has a robust root system (Green, 2011). The profuse root system of hyperaccumulator plants is closely linked to the existence of various types of microbes, which is an important role in the contaminant degradation in rhizosphere (Jiang and Liu, 2010). This microbes will help to breakdown heavy metal compounds to more available form for roots absorption, thus increase the extraction of metals from soil by accumulator plants. Residual heavy metal concentration in soil sown with test plant has been presented on Table 4. As presented, initial soil concentration was 62.5 mg/kg, but residual concentration after plant exposure for 15 weeks was 19.55 mg/kg which represented a remediation efficiency of 68.39%. The most reduced metal concentration in soil was with lead sulhate (4.05 mg/kg), an efficiency of 93.07%. Similarly, total sequestered Pb was highest when lead type in soil was sulphate (58.1 mg/kg) than when other lead anions were present (Table 4). This implied that the concomitant sequestration factor was 92.96%. Generally, sequestration factor ranged from 67.9 – 92.96%. This means that the plant was most likely to sequester the metal than bioaccumulate it in the above ground plant parts.

This is in accordance to a research where the application of heavy metal increased 45% chlorophyll formation and three times more photosynthesis rate in spinach (Kavita*et al*., 2014). However, in the lower concentrations of lead, an exclusion mechanism was indicated. An exclusion mechanism has also been reported by Krämer (2010) in the green alga *Chlorella vulgaris*.

The sequestration efficiency which is the percentage of heavy metals remediated from the soil as well as the foliar concentrated efficiency, increased with decreasing concentration of lead indicating that faster remediation of lead by the plant occurs with decreasing concentration of the metal thus more metal is remediated using the leaves in lower concentration of heavy metal than in higher concentrations.

### Phytoremediation Assessment

Lamhamdi*et al*. (2011) reported that chelation therapy binds the metal and allows the removal of excess or toxic metal from the system rendering it immediately non-toxic and reducing the late effects. Murali*et al*. (2014) reported that the use of sodium acetate as chelating agent gave the highest percentage removal of heavy metals in *Pernaviridis*.

### Microbial Count Determination

From this study, it was observed that there was an increase of bacteria and fungi species in the metal exposed plants of the bulk and rhizospheric soil than in the control. However, the predominant bacteria in the bulk and rhizospheric soil was*Bacillus subtilis* while the predominant fungi was *Aspergillus niger*in the control and metal exposed plants. The microorganisms identified in the bulk and rhizospheric soils have been confirmed by Nazarian*et al*. (2016) as active members of bioremediation microbial consortia. Also Omoregbee*et al*. (2015) stated that plant bacteria interaction could stimulate the production of compounds that could alter soil chemical properties in the rhizosphere and enhance heavy metal accumulation in plants.

## Conclusion

Heavy metals such as Pb can be taken up by roots into the plant tissues through soil. Pb uptake by *Eleusine indica* had no damaging effects on the growth and metabolic activities of the plant. This could be based on the fact that *E. indica* has phytoremediative capacity as it can tolerate a range of abiotic stressors. The present study revealed that *E. indica* has significant contaminant capacity for Pb^2+^ irrespective of the anion combination. However, Pb^2+^ from lead (11) nitrate caused comparatively higher stress - mediated effects on *E. indica*. This study suggest that metabolic pathways and mechanisms of tolerating pb^2+^ of different origin of plant. This misunderstanding is a study for another time.

## Acknowledgments

The authors appreciate Pascal Okoye for his assistance during the study. The study was privately funded.

